# Optimal inhibitory-to-excitatory ratio governs slow and fast oscillations for enhanced neural communication

**DOI:** 10.1101/2025.06.22.660973

**Authors:** Jung Young Kim, Sang Wan Lee, Demian Battaglia, Jee Hyun Choi, Soon-Hyung Yook

**Affiliations:** Department of Bio and Brain Engineering, Korea Advanced Institute of Science and Technology (KAIST), Daejeon, Korea; Computational Cognitive & Systems Neuroscience Laboratory, Brain Science Institute, Korea Institute of Science and Technology (KIST), Seoul, Korea; Department of Physics and Research Institute for Basic Sciences, Kyung Hee University, Seoul, Korea; Department of Brain and Cognitive Sciences (KAIST), Daejeon, Korea; Kim Jaechul Graduate School of AI (KAIST), Daejeon, Korea; Center for Neuroscience-inspired AI (KAIST), Daejeon, Korea; INS - Institut de Neurosciences des Systèmes; USIAS - Institut d’Etudes Avancées de l’Université de Strasbourg - Institute for Advanced Study; LNCA - Laboratoire de neurosciences cognitives et adaptatives; Division of Bio-Medical Science & Technology, Korea University of Science and Technology, Daejeon, Korea

**Keywords:** neural oscillations, synchronization, E/I ratio, neural computation

## Abstract

Neural oscillations at distinct frequency bands facilitate communication within and between neural populations. While single-frequency oscillations are well-characterized, the simultaneous emergence of slow (beta) and fast (gamma) oscillations within the same network remains unclear. Here, we demonstrate that multi-frequency oscillations naturally arise when the ratio of inhibitory-to-excitatory synaptic strength falls within a specific regime using a biologically plausible Izhikevich model. We show that this regime maximizes both information capacity and transmission efficiency, suggesting an optimal balance for neural communication. Deviations from this range lead to single-frequency oscillations and reduced communication efficiency, mirroring disruptions observed in neurological disorders. These findings provide mechanistic insight into how the brain leverages multiple oscillatory frequencies for efficient information processing and suggest a potential biomarker for impaired neural communication.

**SIGNIFICANCE STATEMENT:** Beta (slow) and gamma (fast) oscillations often coexist in the brain, yet their origin and functional role remain unclear. Our study reveals that the inhibitory-to-excitatory synaptic strength ratio governs the emergence of this multifrequency state. Furthermore, we demonstrate that information capacity and transmission efficiency are maximized in this regime, leading to significantly enhanced neural communication. These findings provide mechanistic insight into how multiple oscillatory frequencies support efficient brain function and offer a potential framework for understanding disruptions in neural communication associated with neurological disorders.

## 2. INTRODUCTION

Neural activities arising from the interactions of interconnected neural populations exhibit many intriguing collective phenomena. Among these phenomena, neural oscillations, characterized by rhythmic fluctuations, are particularly important because they organize individual neural activities (K. D. Harris (2005)) and facilitate communication (P. Fries (2005, 2015); T. Akam & D. M. Kullmann (2014)) between populations, which are essential for cognitive functions. Different frequency oscillations are related to specific roles in cognitive functions. Slow oscillations, such as alpha/beta oscillations (15-30 Hz), are often associated with sensory integration and temporal prediction (B. E. Kilavik et al. (2013); H. Tan et al. (2016); M. Lundqvist et al. (2024)). Fast oscillations (gamma oscillation, 40-120 Hz) play different roles, including attention, perception, and information routing (L. L. Colgin et al. (2009); A. Fernandez-Ruiz et al. (2023)). The emergence of each of these frequency oscillation has been explained by the neural circuits composed of excitatory-inhibitory (B. Voloh & T. Womelsdorf (2016); C. Börgers & N. Kopell (2005)) or inhibitory-inhibitory neuronal interactions (X.-J. Wang & G. Buzsáki (1996a); J. A. White et al. (1998)). More recently, several studies have emphasized the combined effect of distinct frequency oscillations, particularly beta and gamma (slow and fast) oscillations, in higher-order cognitive functions such as working memory (M. Lundqvist et al. (2016, 2018); C. G. Richter et al. (2017)) and predictive coding (A. M. Bastos et al. (2020)). These findings suggest that understanding how slow and fast oscillations simultaneously emerge within a neural population provide deeper insights into their role in cognitive functions.

Computational studies have shown the emergence of slow and fast oscillations using various approaches, including the incorporation of different inhibitory neuron types with specific connection strength (S. Keeley et al. (2017); R. Sanchez-Todo et al. (2023)), and tuning of current input strength and decay time constants in single inhibitory circuits (H. Bi et al. (2020)). While these studies provided valuable insights, they often overlooked the distinct spiking properties of excitatory and inhibitory neurons. For example, excitatory neurons often exhibit regular spiking dynamics (V. Jacob et al. (2012); P. Schwindt & W. Crill (1999); S. Franceschetti et al. (1998)), whereas inhibitory parvalbumin interneurons are characterized by fast-spiking activity (X.-J. Wang & G. Buzsáki (1996b); R. Nitsch et al. (1990)). These intrinsic differences introduce distinct time constants that enable the simultaneous generation of slow and fast oscillations depending on synaptic strength.

Furthermore, changes in synaptic strength between excitatory and inhibitory neurons influence neural communication. Strong excitatory input increases redundancy in neural activity due to highly correlated activities (M. A. Dichter & G. Ayala (1987)) which reduces the number of possible patterns for communication. Conversely, excessive inhibition suppresses neural activity, preventing information transmissions. Thus, maintaining balance between excitation and inhibition ratio (E/I ratio) is critical for optimizing information capacity and transmission, both of which are essential for efficient neural communication (W. L. Shew et al. (2011)). Experimental studies further support this, showing that neurological disorders are often associated with disrupted E/I balance (V. S. Sohal & J. L. Rubenstein (2019); S. B. Nelson & V. Valakh (2015)). These findings suggest the potential relationship between neural communication and the emergence of slow and fast oscillations, both of which depend on excitatory-inhibitory interactions.

In this study, we investigate the origin of the slow and fast frequency oscillatory activity in neural populations. Using the Izhikevich model, which incorporates the distinct properties of excitatory and inhibitory neurons, we find an optimal E/I ratio regime for the coexistence of slow and fast oscillations. We also show that information capacity and transmission efficiency are maximized in the same regime. These findings provide clear evidence that efficient neural communication is closely related to the multi-frequency oscillation.

## 3. MATERIALS AND METHODS

### 3.1. Model and Simulation

We construct the neural networks with *N* neurons, composed of *N*_*E*_ excitatory and *N*_*I*_ inhibitory neurons. Based on physiological evidence (S. Marom & G. Shahaf (2002)), we set *N*_*E*_ = 4*N*_*I*_ .

To simulate the neural dynamics, we use the Izhikevich model (E. M. Izhikevich (2003)) described as follows:

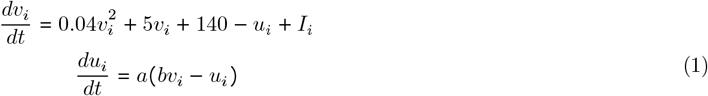

where *v*_*i*_ and *u*_*i*_ represent the membrane voltage and recovery variable of *i*-th neuron, respectively, and *I*_*i*_ is the total incoming current. If *v*_*i*_ exceeds 30 mV at 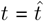, it is reset to *v*_*i*_ ← *c* and updates *u*_*i*_ as *u*_*i*_ ← *u*_*i*_ + *d*. The time 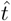 is recorded as the spike time. The dimensionless parameters *a, b, c*, and *d* govern the spiking dynamics.

The total incoming current *I*_*i*_ is given by

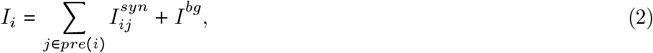

where *pre*(*i*) is the set of presynaptic neurons connected to *i*, 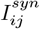 is the synaptic current from presynaptic neuron *j* to *i*, and *I*^*bg*^ denotes the background synaptic current.

The synaptic current 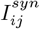 is expressed as

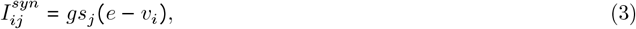

where *g* is the synaptic strength of presynaptic neurons with different values for excitatory and inhibitory neurons, and *s*_*j*_ represents the synaptic activity. The equilibrium potential *e* is set to *e* = 0 mV for excitatory presynaptic neurons and *e* = −80 mV for inhibitory presynaptic neurons.

The synaptic activity *s*_*j*_ evolves over time as,

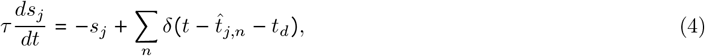

where *δ* (*t*) is the Dirac delta function, and *τ* is the decay time constant. 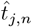 denotes the *n*-th spike time of neuron *j*, and *t*_*d*_ is the synaptic delay. We use *t*_*d*_ = 1 in the following simulations.

Each pair of neurons is connected with a probability *p*_*xy*_ where *x, y* (∈ {*E, I*}) denote the presynaptic (*x*) and postsynaptic (*y*) neuron types. For simplicity, we assume the same connection probability regardless of postsynaptic neuron type *y*, i.e., *p*_*EE*_ = *p*_*EI*_ = *p*_*E*_ and *p*_*IE*_ = *p*_*II*_ = *p*_*I*_ .

The background synaptic current *I*^*bg*^ is generated by external inputs with firing rates following a Poisson distribution with rate *v*_*ext*_ = 2 kHz. Each spike from these neurons induces synaptic activity, described in Eq. 4 with *τ* = 5 ms and *t*_*d*_ = 0 ms, and provides synaptic current satisfying Eq. 3 with *g* = 0.001. Each external input source is connected to each neuron with a probability of *p* = 0.1.

To reflect the distinct physiological properties of excitatory and inhibitory neurons, we use different parameters for each type. For excitatory neurons, we use *a* = 0.02, *b* = 0.2, *c* = −65, and *d* = 8 in Eq. 1 to produce regular spiking dynamics (E. M. Izhikevich (2003)). These parameters are characteristic of cortical excitatory neurons (V. Jacob et al. (2012); P. Schwindt & W. Crill (1999); S. Franceschetti et al. (1998)). The excitatory neurons are connected to the other neuron with probability *p*_*E*_ = 0.2 (S. Mensi et al. (2012); S. P. Brown & S. Hestrin (2009)). We set the synaptic decay time constant as *τ* = 5 ms. For inhibitory neurons, we use *a* = 0.02, *b* = 0.25, *c* = −65, and *d* = 2, which generate fast spiking dynamics as observed in parvalbumin-positive interneuron (X.-J. Wang & G. Buzsáki (1996b); R. Nitsch et al. (1990)). The connection probability is set to *p*_*I*_ = 0.3 reflecting the denser projection of inhibitory interneurons. The synaptic decay time is set to *τ* = 6 ms, which is slower than that of excitatory synapses (D. Curtis & J. Eccles (1959); A. J. Smith et al. (2000)).

For numerical simulations, we use the 4th order Runge-Kutta method with time step Δ*t* = 0.01 ms. Data simulated prior to reaching the steady state (*t* < 1 *s*) are excluded from the average.

### 3.2. Coherence and synchrony

The network synchrony is quantified by two complementary parameters: population coherence (*ρ*) and spike train synchrony (*R*). Population coherence (*ρ*) measures the consistency of collective neuronal activity, defined as (M. Di Volo & A. Torcini (2018))

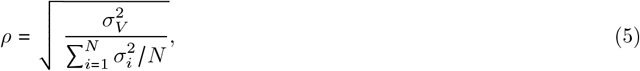

where *σ*_*i*_ and *σ*_*V*_ are the standard deviation of *v*_*i*_(*t*) and 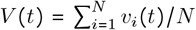, respectively. The coherent activity is associated with a positive value of *ρ*, which ranges from 0 (incoherence) to 1 (perfect coherence).

The spike train synchrony *R* quantifies the consistent temporal alignment of neuronal spikes defined as

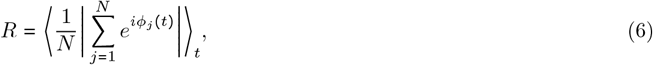

where ∣.∣ and ⟨.⟩_*t*_ denote the modulus of a complex number and its average over time (Y. Kuramoto & Y. Kuramoto (1984)). *ϕ*_*j*_(*t*) is the spike phase of *j*^*th*^ neuron, defined as a linear interpolation between consecutive spikes, treating the interval between two spikes as one cycle. Intervals shorter than 5 ms are disregarded to avoid rapid phase shifts. *R* ranges from 0 (complete desynchronization) to 1 (complete synchronization). These two parameters are computed with 20 randomly selected 1 s window for each simulation, then averaged to obtain the final values.

### 3.3. Network frequency

We characterize the collective oscillatory activity by network frequency *f*_*net*_ (N. Brunel & X.-J. Wang (2003)), determined from the peak positions of Fourier transform of *V* . Multiple values of *f*_*net*_ are obtained if multi-frequency oscillatory activity exist, but integer multiples of other frequencies are excluded to remove harmonics.

### 3.4. Information capacity and transmission efficiency

We first define the spike state vector set *X* = {*X*_1_, *X*_2_, …, *X*_*T*_}, which transforms neural activity into information quantity. The spike time of excitatory neuron is binned into 2 ms intervals and transformed into a binary vector 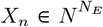, where (*X*_*n*_)_*i*_ = 1 if the neuron *i* is active in the *n*-th bin, and (*X*_*n*_) _*i*_ = 0 otherwise, as described in Fig. 4(a).

The information capacity *Ĥ* is calculated by Renyi’s *α*-order entropy with matrix-based kernel estimator *S*_*α*_ (L. G. S. Giraldo et al. (2014)):

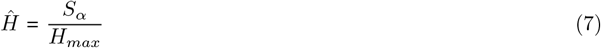

where *H*_*max*_ is the maximum Shannon information entropy, ensuring that *Ĥ* is normalized between 0 and 1.

Renyi’s entropy is particularly suitable for analyzing high-dimensional spike state vectors, and converges to Shannon information entropy as *α* → 1. Here, we set *α* = 1.01 to approximate Shannon information entropy while leveraging the advantages of Renyi’s entropy.

*S*_*α*_ is defined as

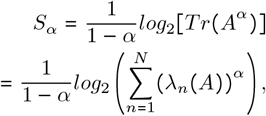

where *Tr* (.) denotes the trace of the matrix, and *Tr* (*A*^*α*^) equals the sum of eigenvalues *λ*_*n*_ of the normalized Gram matrix *A* ∈ *R*^*T* ×*T*^ due to its symmetric property. The elements of *A* are given by

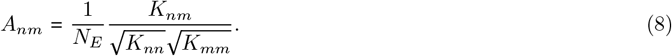

Here, *K* is the Gram matrix *K* ∈ *R*^*T* ×*T*^ defined as

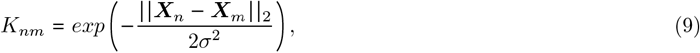

where ∣∣.∣∣ denotes *L*_2_ -norm, and *σ* is the standard deviation of the kernel, set as 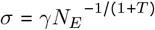 with *γ*=2, following (B. W. Silverman (1998)).

To quantify the mutual information, we generated a set of random *N*_*E*_-dimensional binary codes *Y* = *Y*_1_, *Y*_2_, …, *Y*_*T*_, where *Y*_*i*_ ∈ *Z*_2_. Then they are converted to currents where 1 mA for a binary 1 and -1 mA for a binary 0, and these input currents are updated every 100 ms. We confirmed that *ρ* remains unchanged with current injection, while *R* is affected. However, the overall behavior in each regime remains unchanged (see Fig. S5 in the Supplemental Materials).

The mutual information is also measured through Renyi’s mutual information between *Y* and *X*, defined as

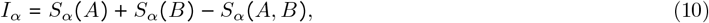

where *S*_*α*_ (*A, B*) is the joint entropy of *A* and *B*, is given by

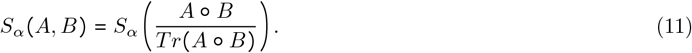

Here, *A* and *B* are the normalized Gram matrices for *X* and *Y*, respectively, and ° denotes element-wise multiplication. Then, mutual information *Î* and transmission efficiency *ϵ* are defined as

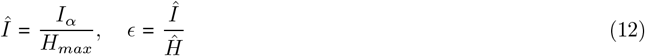

where both *Î* and *ϵ* are bounded between 0 and 1.

### 3.5. Time-lagged cross-correlation

To quantify the temporal relationship between excitatory and inhibitory population activity, we calculate the crosscorrelation *χ* (*τ*) between *v*_*E*_ and *v*_*I*_, defined as

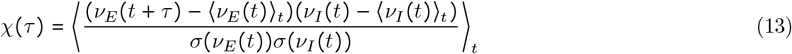

where ⟨.⟩_*t*_ denotes the time average, and *σ*(.) represents the standard deviation. From the obtained *χ*(*τ*), we define *τ*_*EI*_ at which *χ*(*τ*) reaches the maximum

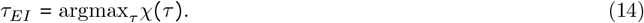

The sign of *τ*_*EI*_ indicates which neuron type leads, a negative *τ*_*EI*_ means that excitatory neurons precede inhibitory neurons, and a positive *τ*_*EI*_ indicates the opposite.

## 4. RESULTS

### 4.1. Oscillatory Regimes

Neural oscillations arise from collective fluctuations of neuronal membrane potentials. To investigate how these fluctuations change as excitatory and inhibitory synaptic strengths (*g*_*E*_ and *g*_*I*_) vary, we first measure the population coherence, *ρ* (for definition, see Materials and Methods).

Fig. 1(a) shows the measured *ρ* for *g*_*E*_ ∈ [0.0, 0.4] and *g*_*I*_ ∈ [0.0, 4.0] on the networks with *N* = 500. From the data, we find that there are low-coherence and high-coherence regimes. The high-coherence regime is observed in both the excitation-dominant and inhibition-dominant areas, separated by the low-coherence regime in the intermediate area. To estimate finite-size effects, we examine the asymptotic behavior of *ρ* by increasing *N* while keeping the in-degree for each neuron and the other model parameters in each regime the same. In the high-coherence regimes, *ρ* converges to values greater than 0.4 as *N* increases (°, ⊲ in Fig. 1(b)). On the other hand, in the low-coherence regime, *ρ* decreases with increasing *N* and approaches 0 (⋆ in Fig. 1(b)). The dashed lines in Fig. 1(a) indicate the boundaries that separate regimes based on the asymptotic behavior of *ρ*.

**Figure 1.**
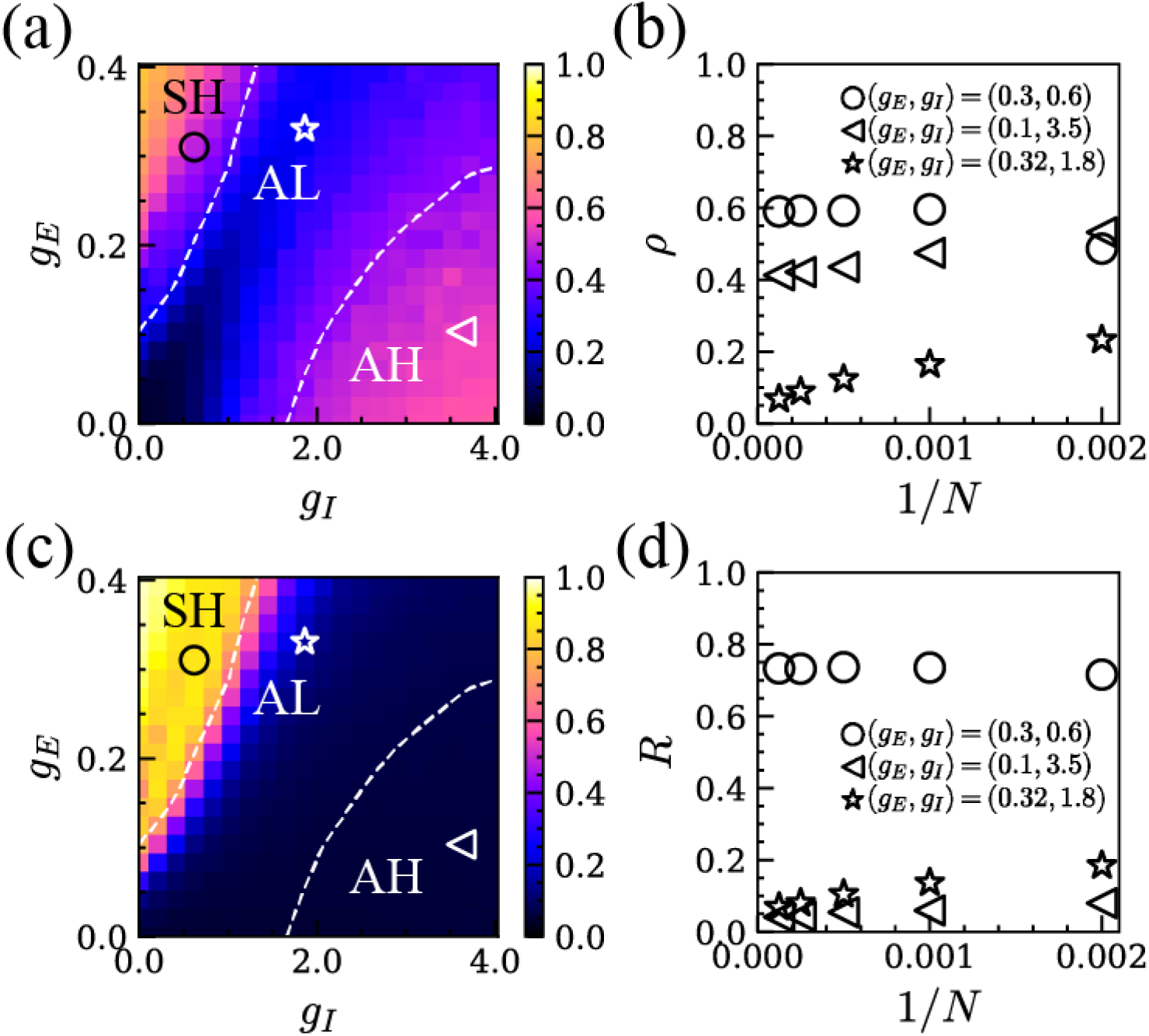
(a) Plot of *ρ*(*g*_*E*_, *g*_*I*_) with *N* = 500. The white dashed line denotes the boundaries that separate the asymptotic behavior of *ρ*. (c) Plot of *R*(*g*_*E*_, *g*_*I*_) with *N* = 500. (d) Plot of *R* against 1/*N* . The symbols in (a) and (c) denote randomly selected values of (*g*_*E*_, *g*_*I*_) from each regime: Circles (°) correspond to (*g*_*E*_, *g*_*I*_) = (0.3, 0.6), triangles (⊲) represent (*g*_*E*_, *g*_*I*_) = (0.1, 3.5), and stars (☆) denote (*g*_*E*_, *g*_*I*_) = (0.32, 1.8), respectively. The regimes are defined as SH (synchronous and high-coherence), AL (asynchronous and low-coherence), and AH (asynchronous and low-coherence).

Collective fluctuations of activity could originate from spike-to-spike synchrony, where the firing of individual neurons triggers the firing of other neurons via excitatory synaptic interactions. It could however also arise in when spike times are not synchronized, where the mean-field activity oscillates, in a way apparent from the monitoring of subthreshold fluctuations (N. Brunel & D. Hansel (2006)). To track whether oscillations are due to spike-to-spike synchrony or asynchronized spike activity, we evaluate as well, in addition to *ρ*, a further measure *R* of spike-to-spike synchrony, defined in Materials and Methods. As shown in Fig. 1(c), three regimes can be distinguished in terms of the joint inspection of parameter-dependent *ρ* and *R* joint variations.

In the excitation dominant area of the (*g*_*I*_ −*g*_*E*_) plane, we observe high *R* and high *ρ*. This regime is thus characterized by spike-to-spike synchrony and we call it the Synchronous High-coherence (SH) regime. In the inhibition dominant area, *ρ* is large but *R* small, indicating High-coherence but Asynchronized spike activity (AH). Between these two regimes, we find a transition zone where not only asynchronized spike activity, as denoted by lower *R*, but also the collective fluctuations is lowered, as shown by low *ρ*. As we will show, however, this Asynchronous Low synchrony (AL) regime transiently exhibits properly more similar to either the AH or the SH regimes. Notably, these three regimes persist regardless of the background input current level (See Fig. S1 in Supplemental Materials).

Fig. 2 illustrate examples of time series and spectrograms in each regime. Each panel shows the membrane potentials of two randomly selected excitatory neurons (*v*_*i*_) and the corresponding power spectrogram of *V* . In the SH regime (*g*_*E*_ = 0.3, *g*_*I*_ = 0.6), excitatory neurons exhibit highly synchronized spiking activity with identical spike times, as shown in Fig. 2(a). The corresponding spectrogram in Fig. 2(b) shows strong oscillatory activity concentrated around 15 Hz. Higher frequency peaks correspond to harmonics of this oscillation and do not reflect distinct oscillatory activity. In the AH regime (*g*_*E*_ = 0.1, *g*_*I*_ = 3.5), membrane potentials exhibit coherent activity, i.e., the increasing and decreasing tendencies of the membrane potentials of the two randomly selected neurons are almost identical. However, the positions of the spiking peaks do not align between two neurons. Furthermore, the spiking interval becomes irregular as shown in Fig. 2(c). Thus, the spiking activity becomes asynchronous in this regime. The period of the membrane voltage fluctuation becomes shorter than that in the SH regime, resulting in fast oscillations around 80 Hz as shown in the spectrogram in Fig. 2(d). Fig. 2(e) shows the time evolution of *v*_*i*_’s in the AL regime (*g*_*E*_ = 0.32, *g*_*I*_ = 1.8). The data clearly show that the neural activities of the two randomly chosen neurons are neither coherent nor synchronous.

**Figure 2.**
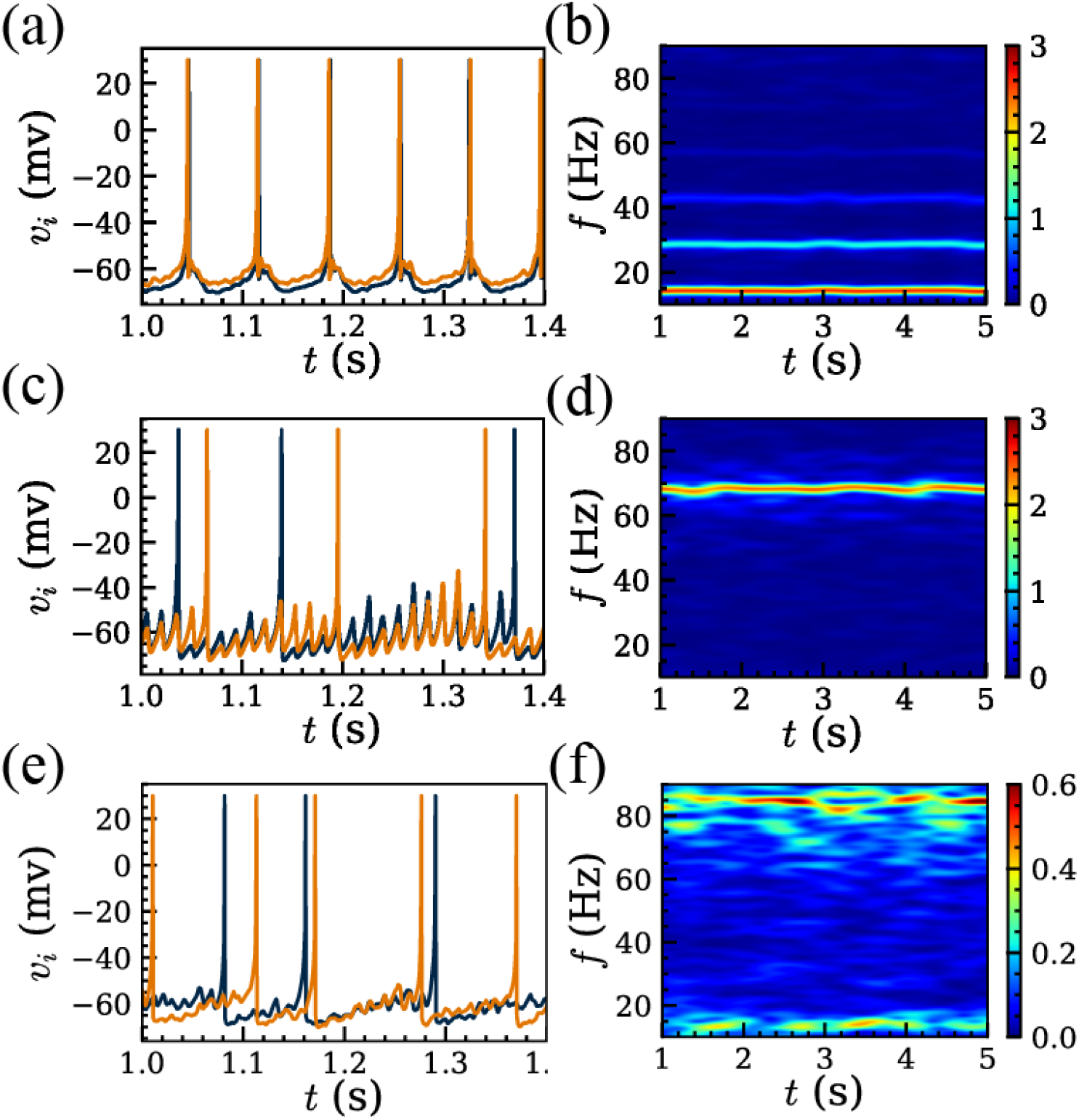
The membrane potentials with time from two randomly selected excitatory neurons (a) and the power spectrogram of *V* (b) at (*g*_*E*_, *g*_*I*_) = (0.3, 0.6) in SH regime. (c) and (d) are the same as (a) and (b) with (*g*_*E*_, *g*_*I*_) = (0.1, 3.5) in AH regime, and (*g*_*E*_, *g*_*I*_) = (0.32, 1.8) in AL regime for (e) and (f).

We find that two distinct frequencies are highlighted in the spectrogram of Fig. 2(f), even though they exhibit weak and fluctuating power. The slow oscillation around 15 Hz originates from the spiking activities with a relatively large period. On the other hand, the fast oscillation around 80 Hz arises from the fast subthreshold oscillations in Fig. 2(e). This is further supported by the conditional probability of *R* and *ρ* to slow and fast oscillation powers (see Fig. S2 in the Supplemental Materials). *R* increases only when slow oscillation power increases, while *ρ* increases as either slow or fast oscillation powers increase. Therefore, in the AL regime, slow and fast oscillations coexist, each originated from distinct regimes.

In Fig. 3(a) we show the network frequency (details in Materials and Methods), *f*_*net*_, measured in AL regime (N. Brunel & X.-J. Wang (2003)). Fig. 3(b) shows *f*_*net*_ against *g*_*I*_ / *g*_*E*_ for a fixed *g*_*E*_ = 0.25, which spans all three regimes. The overall behavior was not affected by the value of *g*_*E*_ (see Fig. S3 in the Supplemental Materials). When *g*_*I*_ / *g*_*E*_ < 4.5, which corresponds to the SH regime, the obtained *f*_*net*_ is approximately 15 Hz. As *g*_*I*_ / *g*_*E*_ increases, a fast oscillation (*f*_*net*_ ≳ 80 Hz) emerges alongside the slow oscillation, and this coexistence persists through the AL regime (4.5 ≤ *g*_*I*_ / *g*_*E*_ < 6.5). Notably, these slow and fast oscillation frequencies do not follow an integer multiple relationship, indicating that the fast oscillation is not merely a harmonic of slow oscillation (see Fig. S4 in the Supplemental Materials). In the AH regime (*g*_*I*_ / *g*_*E*_ ≥ 6.5), the slow oscillation disappears, leaving only the fast oscillation. These results are consistent with the spectrograms in Fig. 2, and show that slow and fast oscillations coexist in the AL regime.

**Figure 3.**
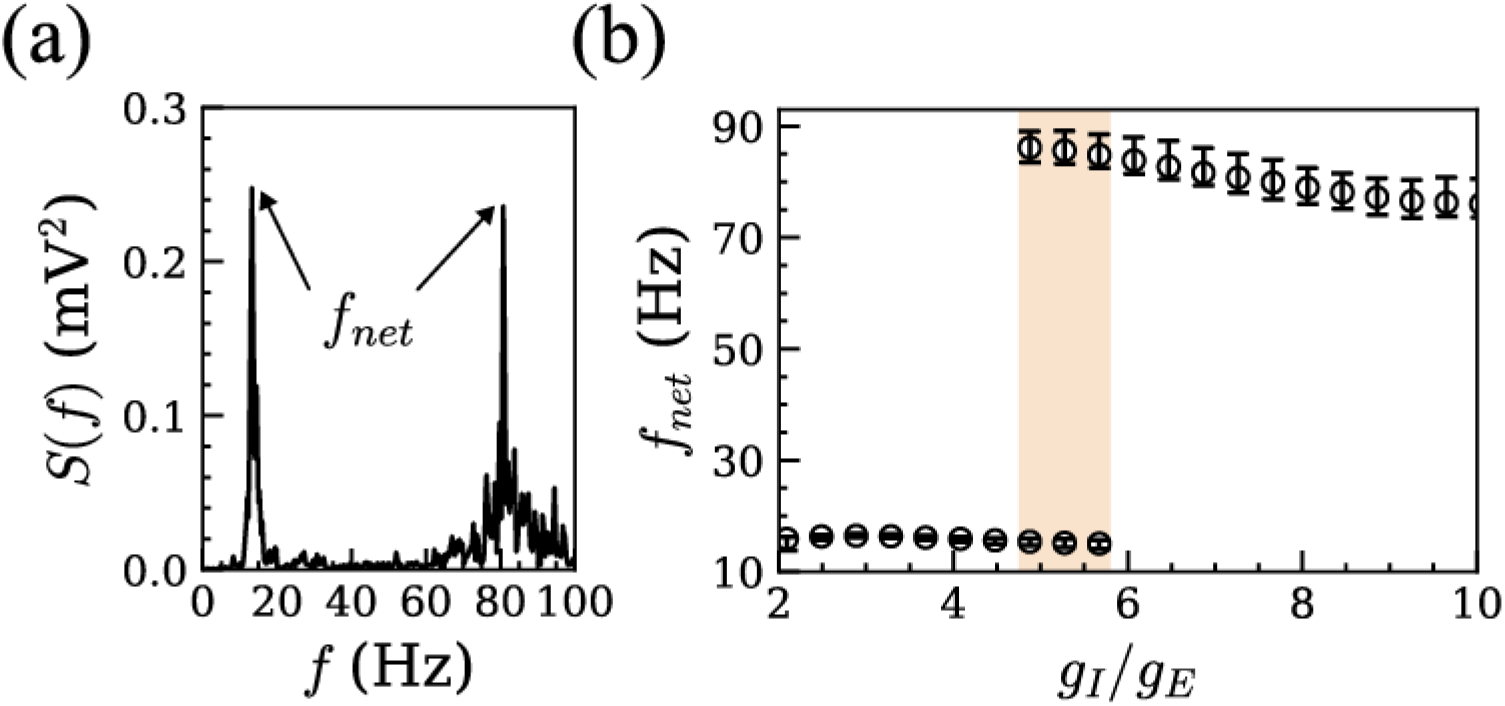
Power spectrum *S*(*f*) of *V* at (*g*_*E*_, *g*_*I*_) = (0.25, 1.32) (a). The arrow indicates two values of *f*_*net*_. (b) *f*_*net*_ against *g*_*I*_ /*g*_*E*_ with *g*_*E*_ = 0.25, which cross three different regimes. Within the AL regime in 4.5 ≤ *g*_*I*_ /*g*_*E*_ < 6.5, two distinct frequency peaks are observed, indicated by the shaded area.

### 4.2. Information Capacity and Transmission Efficiency

From Fig. 1 and Fig. 2, we observe that the characteristic feature of neural activity changes from SH regime to AH regime through AL regime as *g*_*I*_ / *g*_*E*_ increases. Since neural communication and various neurological disorders are known to be closely related to the E/I balance (V. S. Sohal & J. L. Rubenstein (2019); S. B. Nelson & V. Valakh (2015)), it is important to investigate how the ratio *g*_*I*_ / *g*_*E*_ affects information capacity and transmission efficiency.

To estimate the information capacity of the network, we calculate the entropy, *Ĥ*, of spike activity patterns, reflecting the diversity of neural activity (see definition in Materials and Methods). Limited diversity of neural activities constrains the ability to encode upstream input, reducing the amount of information transmitted to downstream populations. Fig. 4(a) illustrates how spike patterns were converted into spike-state vectors for entropy calculation. As shown in Fig. 4(b), *Ĥ* increases as *g*_*I*_ /*g*_*E*_ grows from low values in the SH regime, where synchrony dominates and pattern diversity is limited. In the AL regime, where both synchrony and coherence vanish, *Ĥ* reaches its peak, indicating decreases and the neural activity goes into the AH maximal diversity of activity patterns. Beyond this point, as *Ĥ* regime, where coherence increases again. The behavior of *Ĥ* shows that the AL regime, characterized by the absence of synchrony and coherence, provides the highest capacity for information storage and encoding.

**Figure 4.**
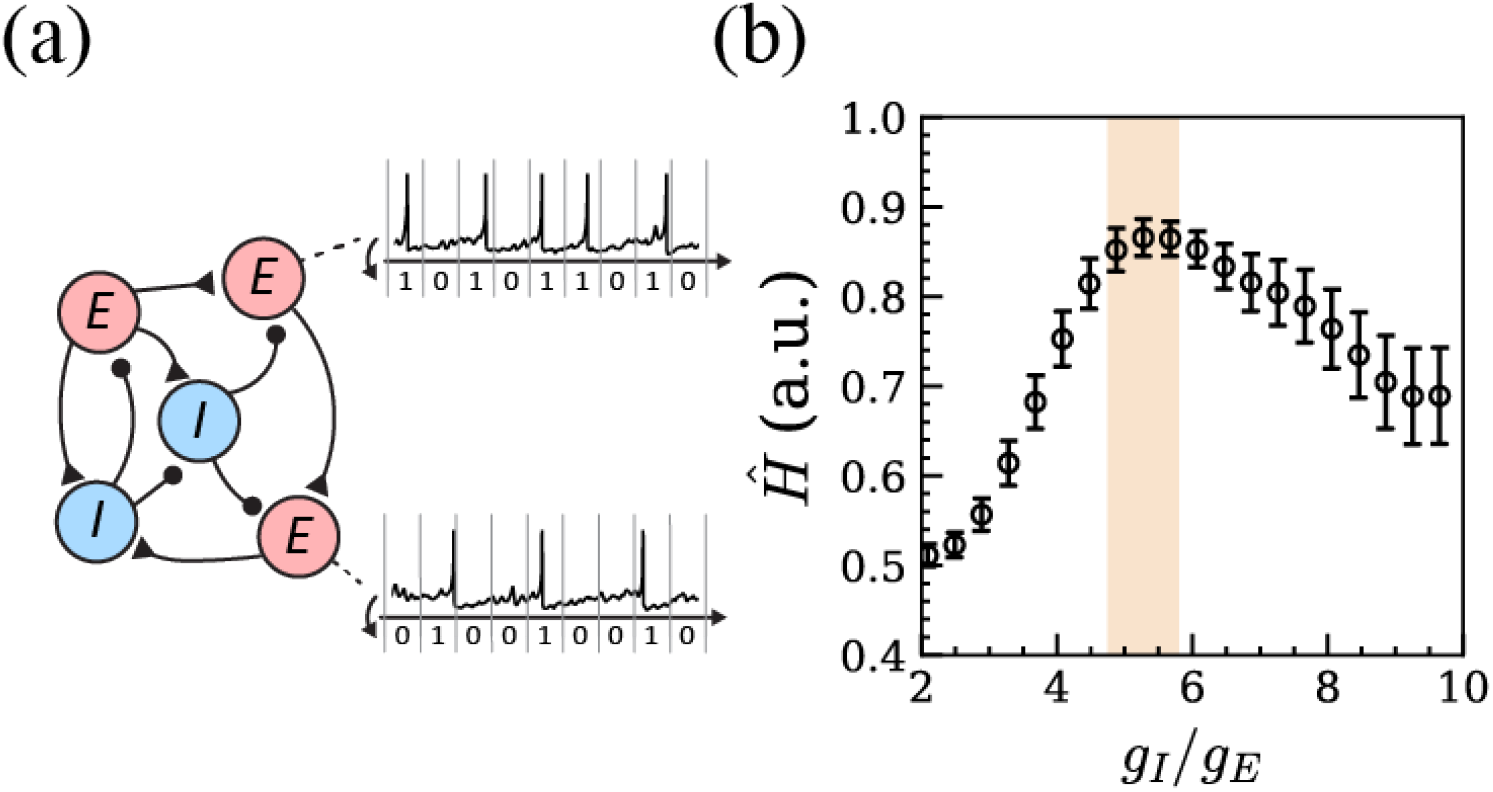
The schematic diagram of converting spike activity into spike-state vector (a) and the normalized information entropy *Ĥ* as a function of the ratio *g*_*I*_ / *g*_*E*_ with *g*_*E*_ = 0.25 (b). The shaded area indicates the regions where two distinct frequency peaks are observed in Fig. 3.

For effective communication, diversity of neural activity patterns alone is insufficient; the input signals must also be accurately encoded into neural responses. To find the effect of E/I balance on the information encoding, we computed the mutual information, *Î*, between input and response activities. Since the total amount of information encoded by the spike activity pattern strongly depends on the value of *g*_*I*_ / *g*_*E*_, we measure the transmission efficiency, *ϵ*, through the normalized mutual information by *Ĥ* (see Materials and Methods).

As shown in Fig. 5(a-b), both *Î* and *ϵ* peaked in the AL regime, where we find that *Ĥ* reaches the maximum (see Fig. 4(b)). Note that the value of *g*_*I*_ /*g*_*E*_ at which *ϵ* reaches its maximum coincides with the value of *g*_*I*_ /*g*_*E*_ at which both *Î* and *Ĥ* also reach their maximum. This indicates that the increase of *Î* is not simply a result of enhanced information capacity but reflects an enhanced encoding process that maximizes the use of available capacity for information transmission.

**Figure 5.**
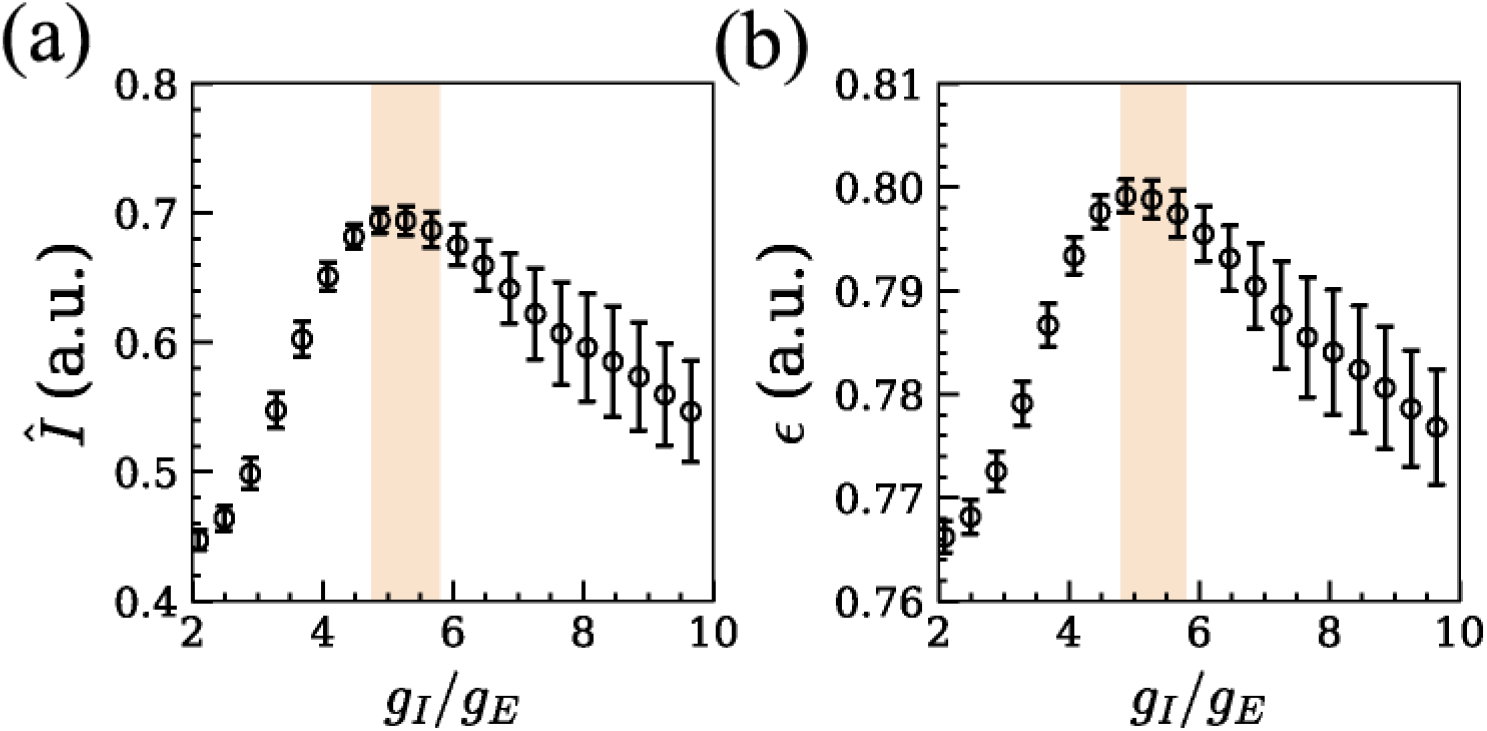
The normalized mutual information *Î* (a) and transmission efficiency *ϵ* (b) as a function of the ratio *g*_*I*_ / *g*_*E*_ with *g*_*E*_ = 0.25. The shaded area indicates the regions where two distinct frequency peaks are observed in Fig. 3.

### 4.3. Temporal relationships between excitatory and inhibitory population activities

Neural oscillations, particularly in the gamma band, are explained by two different dynamical mechanisms. The pyramidal interneuronal network gamma (PING) model explains how oscillations arise from excitatory neurons activating inhibitory neurons (M. A. Whittington et al. (1995))). On the other hand, the interneural network gamma (ING) model (X.-J. Wang & G. Buzsáki (1996a)) explains neural oscillations through mutual inhibition which acts as the pacemaker. To characterize the dynamical properties in each regime, we analyze the temporal relationships between excitatory (*v*_*E*_) and inhibitory (*v*_*I*_) instantaneous firing rates, which represent the number of spikes per unit time. In Fig. 6(a), we present the measured *v*_*E*_ and *v*_*I*_ in the SH regime. The data shows that *v*_*E*_ consistently precedes *v*_*I*_, indicating the behavior predicted by the PING model. On the other hand, in the AH regime, *v*_*I*_ leads *v*_*E*_, reflecting the dynamics of the ING model (Fig. 6(b)).

**Figure 6.**
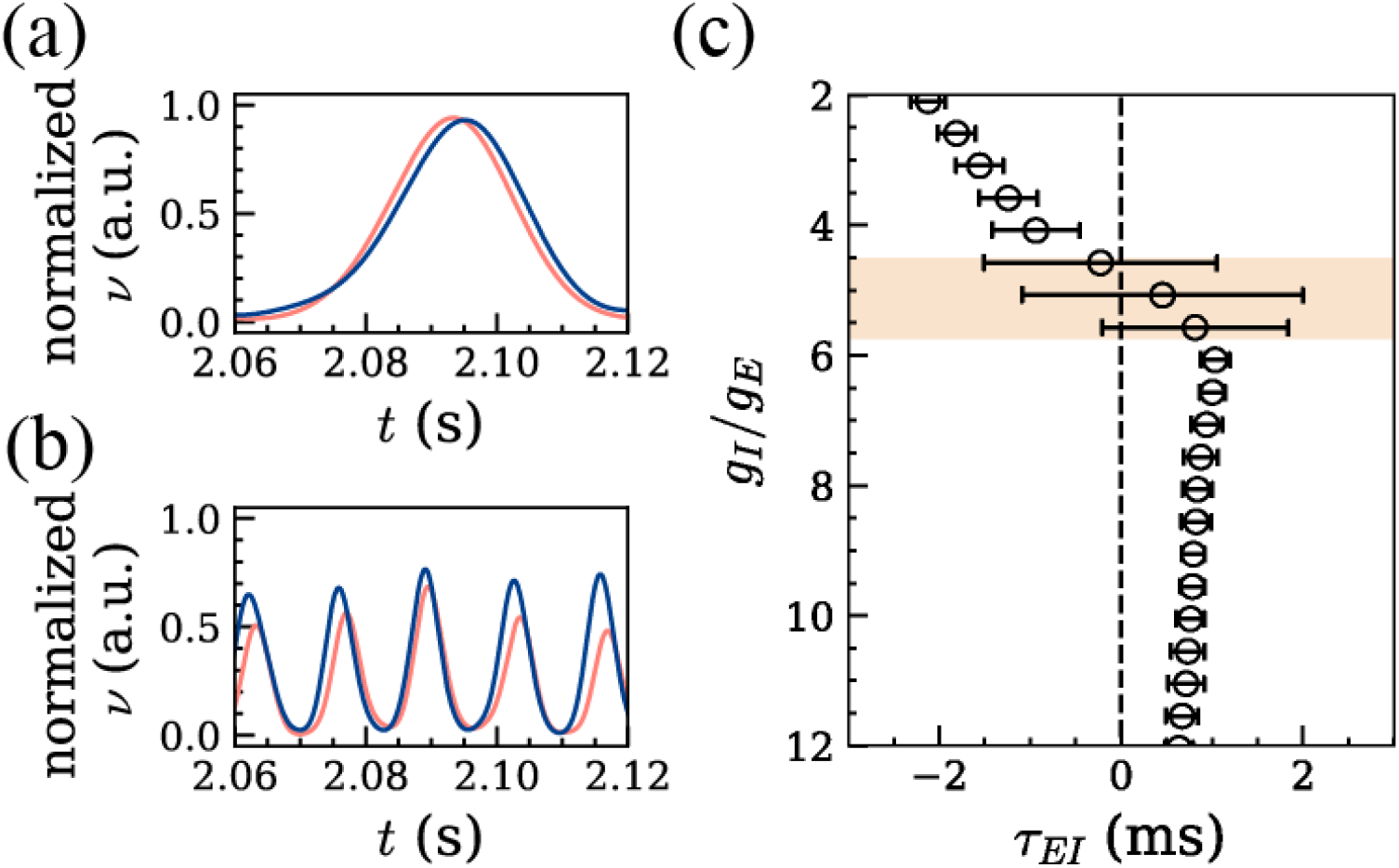
(color online) The example instantaneous firing rate of excitatory (*v*_*E*_, red) and inhibitory (*v*_*I*_, blue) populations when (*g*_*E*_, *g*_*I*_) = (0.25, 0.72) (a) and (*g*_*E*_, *g*_*I*_) = (0.25, 2.21) (b). (c) *τ*_*EI*_ against *g*_*I*_ /*g*_*E*_ when *g*_*E*_ = 0.25, and shaded area indicates the regions where two distinct frequency peaks are observed in Fig. 3. Within this area, two populations does not exhibit a significant preceding relationship.

For a quantitative analysis of these temporal relationships we calculate the time-lagged cross-correlation, *χ* (*τ*), between *v*_*E*_ and *v*_*I*_. Then the time-lag *τ*_*EI*_, at which *χ*(*τ*) becomes the maximum, is estimated (see details in Materials and Methods). Fig. 6(c) shows the relationship between *g*_*I*_ /*g*_*E*_ and *τ*_*EI*_ . In the SH regime, which corresponds to low *g*_*I*_ /*g*_*E*_ values, *τ*_*EI*_ is negative, indicating that excitatory neurons drive to inhibitory neurons. Thus, the neural activities in the SH regime are characterized by the PING model. In the AH regime, which corresponds to high *g*_*I*_ /*g*_*E*_ values, *τ*_*IE*_ becomes positive. This indicates that the AH regime shows a typical behavior of the ING model. However, in the AL regime, which is in between SH and AH regimes with two distinct frequency oscillations, does not exhibit a significant leading role by either excitatory or inhibitory neurons. This suggests that the AL regime lacks a well-defined temporal relationship between the two neuronal populations.

## 5. DISCUSSION

We investigated the emergence of slow and fast frequency oscillations in neural populations by varying the excitatory and inhibitory synaptic strength ratios, incorporating their distinct properties. Our results reveal that slow and fast oscillations emerge in excitatory and inhibitory dominant regimes, respectively. Between these regimes, both oscillations coexist, coinciding with increased information capacity and transmission efficiency. These findings suggest that a multi-frequency oscillatory state is a hallmark of optimal neural communication. On the other hand, shifts to single-frequency oscillations by excessive excitation or inhibition indicate the disrupted communication state, potentially impairing network function.

Neural oscillations have traditionally been explained by two classes of models, both supported by pharmacological manipulation of brain tissue. The PING model accounts for gamma oscillations as excitatory-driven rhythms emerging from excitatory-inhibitory interactions (M. A. Whittington et al. (1995); P. Tiesinga & T. J. Sejnowski (2009)), while the ING model describes inhibitory-driven oscillations generated through mutual inhibition X.-J. Wang & G. Buzsáki (1996a); J. A. White et al. (1998). Our findings reveal that under strong excitatory or inhibitory synaptic input, neural dynamics align with these classical models. However, neither PING nor ING models account for the emergence of multi-frequency oscillations within the same network.

To explain this phenomenon, several studies have investigated conditions that give rise to multi-frequency oscillations within excitatory-inhibitory populations. Brunel identified a specific E/I region where two distinct frequency oscillations coexist, similar to our findings (N. Brunel (2000)). However, this region exists only when input strength is just above the firing threshold and disappears as input strength increases. In contrast, our findings show that the AL regime expands with stronger inputs, suggesting a more robust mechanism. Similarly, Bi et al. demonstrated that slow and fast oscillations can emerge in a balanced excitatory-inhibitory network, in the fluctuation-driven oscillation regime (H. Bi et al. (2021)). Their mean-field solution exhibits a stable focus, which may explain the emergence of multifrequency oscillatory activity, though this regime shows low coherence in our results. However, the model predicts that fast oscillation frequency is integer multiples of slow oscillation frequency, which is not necessary for experimentally observed beta and gamma oscillations (M. Lundqvist et al. (2016); C. G. Richter et al. (2017)). Our model does not exhibit this harmonic relationship, further emphasizing the need for alternative mechanisms. Together, these findings suggest that previous models are insufficient to explain simultaneous beta and gamma oscillations generation. This highlights the importance of incorporating distinct excitatory and inhibitory neuron properties to better capture the complexity of multi-frequency oscillations in neural populations.

Beyond excitatory-inhibitory population, inhibitory neuron subtypes may also play a role in the co-occurrence of slow and fast oscillations. For example, Chen et al. demonstrated that beta and gamma oscillations in visual cortex are modulated by parvalbumin (PV) interneuron and somatostatin (SOM) interneuron, respectively, by optogenetic manipulation (G. Chen et al. (2017)). In addition, Keeley et al. proposed that mutual inhibition between two inhibitory subtypes with distinct synaptic decay time constants can generate slow and fast oscillations (S. Keeley et al. (2017)). However, inhibitory neuron subtypes, especially PV interneuron and SOM interneurons, differ not only in synaptic properties but also in intrinsic spiking dynamics (M. Nassar et al. (2015)) and network structure (S. B. Nelson (2002)). This suggests that their different contributions to oscillatory activity extend beyond simple synaptic interactions. Thus, incorporating these distinct inhibitory neuron properties into the model could reveal more complex oscillatory dynamics, providing a comprehensive framework for understanding cortical oscillation activities beyond slow and fast oscillation dynamics.

We also explored the role of multi-frequency oscillations in neural network function by analyzing the diversity of spike activity patterns (entropy) and mutual information between input and response activities. While the definition of neural information is still debated, information-theoretic approaches provide valuable insights into neural coding. Importantly, experimental evidence supports the relevance of entropy-based measures in neural systems. Several studies have shown that neural populations operate near maximal information entropy, minimizing redundant activity across neurons (S. Laughlin (1981); E. P. Simoncelli & B. A. Olshausen (2001)). Although prior studies did not explicitly link these measures to oscillatory dynamics, Shew et al. have demonstrated that both entropy and mutual information are maximized within a balanced E/I regime through multielectrode array recordings (W. L. Shew et al. (2011)). Consistent with this, Gireesh and Plenz observed multi-frequency oscillatory activity emerges under the network condition similar to Shew et al.’s work (E. D. Gireesh & D. Plenz (2008)). These findings provide an empirical foundation for using entropy as a proxy for tracking the information processing of neural networks and its relationship with multi-frequency oscillatory activity.

It is notable that the specific E/I ratio values support optimal information transmission efficiency, along with the multi-frequency oscillation in this regime. This suggests that deviations from this ratio could significantly disrupt neural communication. Consistently, many neurological disorders, including epilepsy, schizophrenia, and autism spectrum disorder (ASD), are associated with disruptions in E/I balance (V. S. Sohal & J. L. Rubenstein (2019); S. B. Nelson & V. Valakh (2015)). Such disorders often involve abnormal oscillatory patterns in broad range of frequencies, rather than simple changes in oscillatory power. For example, Parkinson’s disease (PD) patients frequently exhibit elevated beta oscillations and reduced gamma activity (M. Weinberger et al. (2009)), whereas Alzheimer’s disease (AD) patients display the opposite trend, with increased gamma power and weakened beta oscillations (J. Wang et al. (2017)). These results suggest that neural computation is not determined by the enhancement or suppression of a specific frequency but rather by maintaining balanced multi-frequency oscillation state, which aligns with our findings. This perspective challenges the conventional view that abnormal oscillatory patterns in neurological disorders originate solely from shifts in single-frequency power, instead highlighting the importance of maintaining a balanced and dynamic multi-frequency state for healthy neural function.

While our study provides valuable insights into how multi-frequency oscillations emerge and contribute to neural communication, it has certain limitations. First, our model assumes homogeneous neuronal properties for each type neuron, whereas real cortical circuit contain diverse excitatory and inhibitory subtypes with distinct spiking dynamics and connectivity patterns (S. B. Nelson (2002)). Second, our model does not account for spatial network organization, which is known to shape neuronal synchronization. Future work could explore how spatial structure influences the emergence and function of multi-frequency oscillatory state. Additionally, integrating these principles into whole-brain models could further our understanding of large-scale neural computations, with potential applications in neurological disease modeling.

In summary, we demonstrate that the coexistence of slow and fast frequency oscillations corresponds to an enhanced neural communication state, emerging within a specific E/I ratio regime. This finding establishes a mechanistic link between E/I balance, oscillatory dynamics, and information processing. This link advances our understanding of neural computation with potential clinical applications.

## ACKNOWLEDGMENTS

We thank the members of the Computational Cognitive and Systems Neuroscience Laboratory in KIST and Physics of Complex Systems and Informatics Laboratory in Kyung Hee University for their helpful discussions. This work was supported by the Basic Science Research Programs (RS-2022-NR070539, 2021R1A6A3A01087727) and the Global Cooperative Convergence Research Program (RS-2024-00460958) by National Research Foundation of Korea (NRF), the intramural grant of the Korea Institute of Science and Technology (2E32901). This work was also supported by Basic Science Research Program through the National Research Foundation of Korea (NRF) funded by the Ministry of Education (Republic of Korea) and the Korea government (MSIT) (grant number: NRF-2019R1F1A1058549 and NRF-2022R1F1A1073629).

## AUTHOR CONTRIBUTIONS

J.Y.K. performed the modeling, analysis, visualization, and manuscript writing. S.Y. contributed to the modeling, supervised the research, and manuscript writing. J.H.C. was responsible for conceptualization, supervision, and manuscript writing. S.W.L. and D.B. revised the manuscript. All authors reviewed and approved the final version of the manuscript.

