## Supplementary materials for "Optimal inhibitory-to-excitatory ratio governs slow and fast oscillations for enhanced neural communication"

**This PDF file includes:**

Figure S1
Figure S2
Figure S3
Figure S4
Figure S5

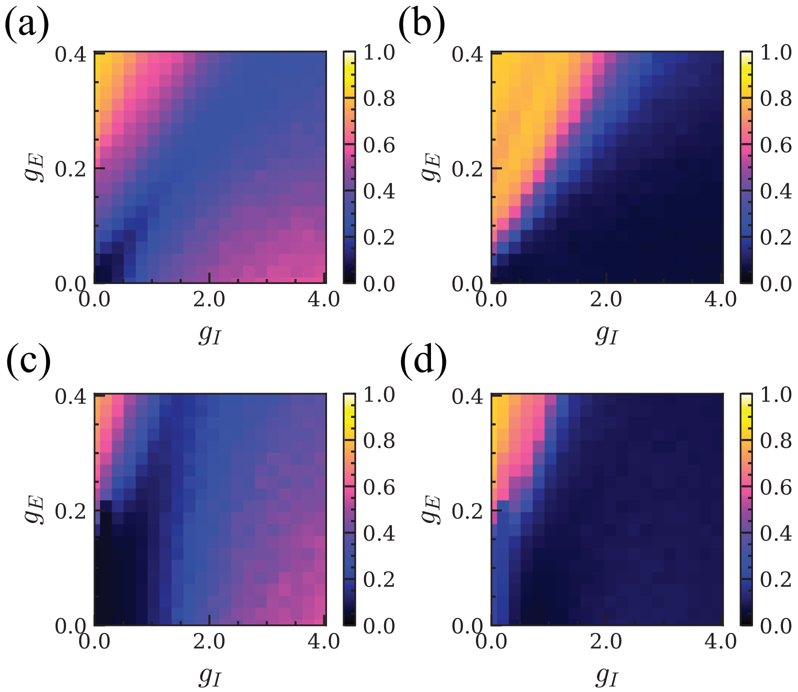


**Figure. S1.** Plots of $\rho$ (a) and $R$ (b) with $N = 500$ under a background neuron firing rate of 1000 Hz. Panels (c) and (d) shows the same but with the firing rate of 4000 Hz. As the background firing rate increases, the average input current to each neuron also increases. The figure shows that the high-coherence and low-coherence regimes are preserved regardless of the input current level, although the I-E ratio for the AL regime increases.

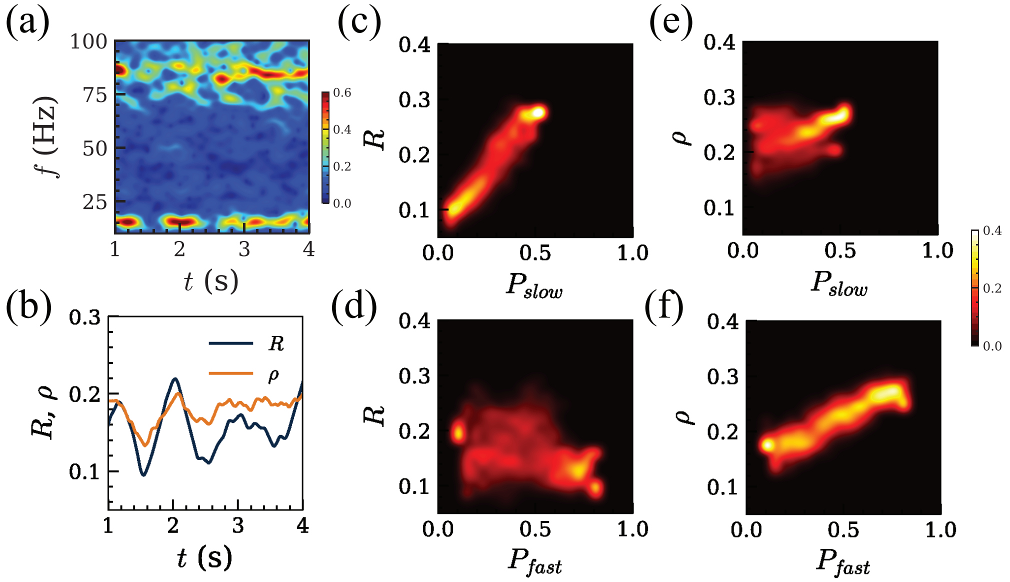


**Figure. S2.** Changes in $R$ and $\rho$ with slow and fast oscillation powers at $g_I/g_E = 5.5$ for $g_E = 0.25$. (a) Power spectrogram shows the fluctuations in slow (10-20 Hz) and fast (80-90 Hz) oscillations, along with the corresponding fluctuations in $R$ and $\rho$ (b). The color scale represents oscillation power. (c-f) Conditional probability distributions $P(R|P_{slow})$ (c), $P(R|P_{fast})$ (d), $P(\rho|P_{slow})$ (f), and $P(\rho|P_{fast})$, where $P_{slow}$ and $P_{fast}$ correspond to the slow (10-20 Hz) and fast (80-90 Hz) oscillation powers, respectively. Here, we computed $R$, $\rho$, and oscillation powers within 1 s windows with 0.05 s time step. The color scale is conditional probability. The figure demonstrates that $R$ increases with stronger slow oscillation, while $\rho$ increases with both frequency oscillations, aligning with the characteristics of SH and AH regimes.

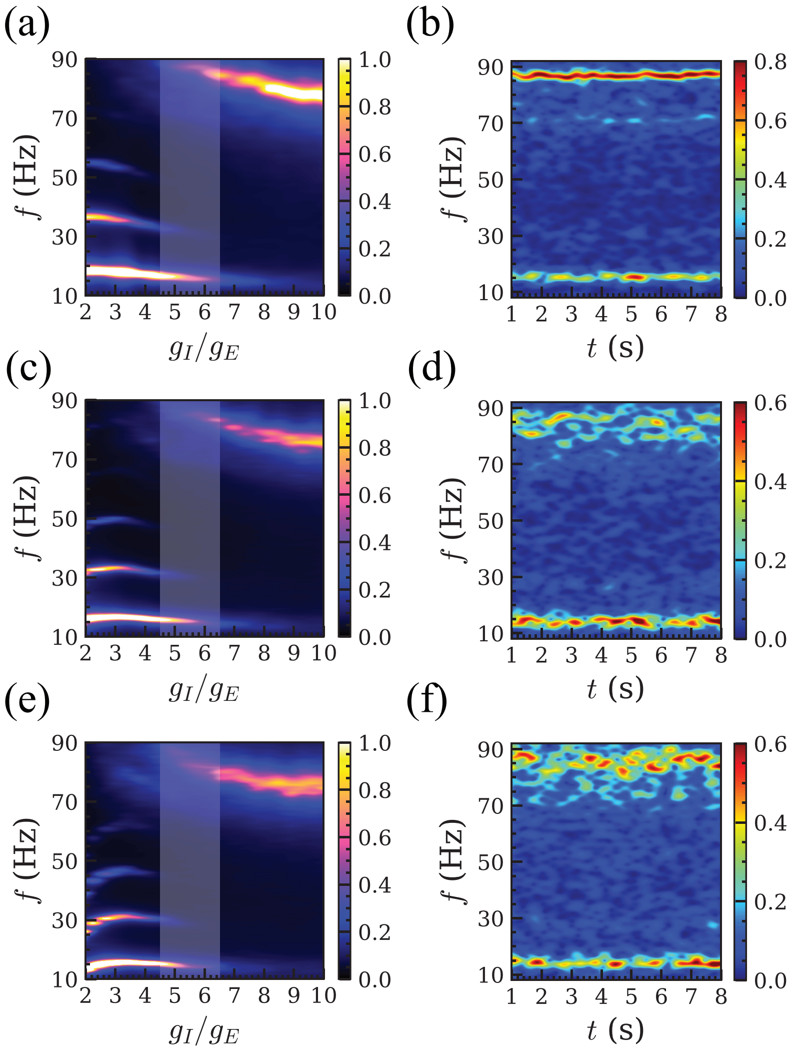


**Figure. S3.** Power spectrum $S(f)$ as a function of $g_I/g_E = 5.57$ in (a), along with the sample power spectrogram at $g_I/g_E = 5.57$ for $g_E = 0.2$ in (b). The color scale represents the power of oscillatory activity, where warmer colors indicate higher values. Panels (c) and (d) shows the same for $g_E = 0.25$, and panels (e) and (f) for $g_E = 0.3$. The white shaded areas in (a),(c), and (e) indicate the regime $4.5 \leq g_I/g_E \leq 6.5$ where two distinct frequency peaks are observed in Fig. 3. The figure demonstrates that multi-frequency oscillatory behavior is preserved over different values of $g_E = 0.2, 0.25,$ and $g_E = 0.3$ with varying $g_E/g_I$.

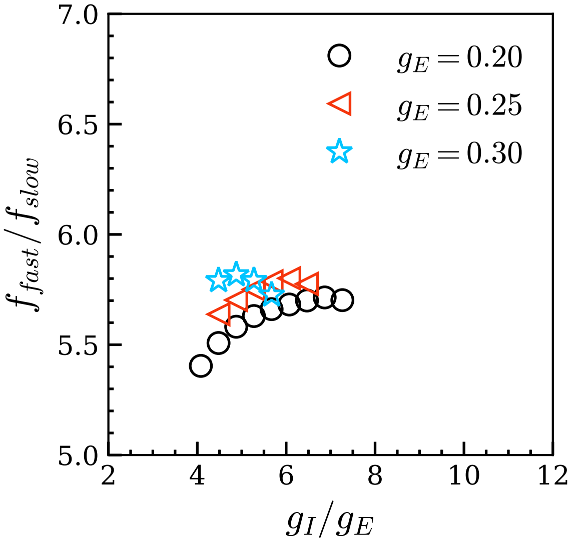


**Figure. S4.** Ratio of fast to slow oscillation frequency ($f_{fast}/f_{slow}$) for different values of $g_E = 0.2,\ 0.25,$ and $0.3$ as a function of $g_I/g_E$. The ratios range between 5.5 and 6.0, indicating that the fast and slow oscillation frequencies do not follow an integer multiple relationship.

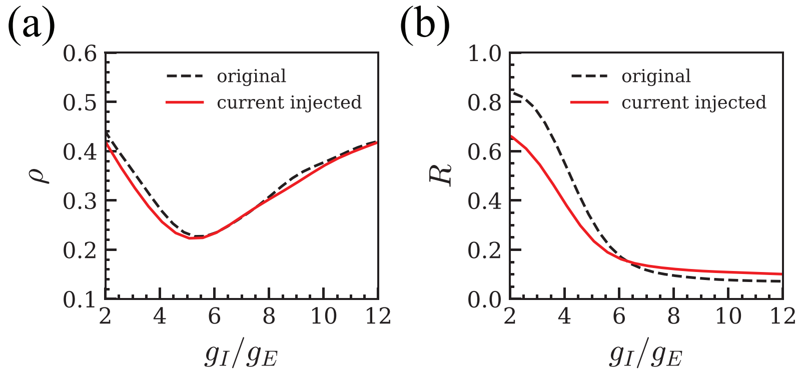


**Figure. S5.** Plots of $\rho$ (a) and $R$ (b) as a function of $g_I/g_E$ at $g_E = 0.25$ comparing without current injection (black dashed line) and with current injection (red line). Although $R$ changes due to perturbed spike timings from current injection, the overall behaviors of each regime is preserved.
